# Optical spike detection and connectivity analysis with a far-red voltage-sensitive fluorophore reveals changes to network connectivity in development and disease

**DOI:** 10.1101/2020.10.09.332270

**Authors:** Alison S. Walker, Benjamin K. Raliski, Kaveh Karbasi, Patrick Zhang, Kate Sanders, Evan W. Miller

**Affiliations:** Departments of Chemistry, University of California, Berkeley, 94720, United States of America; Molecular & Cell Biology, University of California, Berkeley, 94720, United States of America; Helen Wills Neuroscience Institute, University of California, Berkeley, 94720, United States of America

**Author notes:** **Corresponding Author:** Evan W. Miller, B84 Hildebrand Hall, Room 227, University of California, Berkeley, 94720-1460.

## Abstract

The ability to optically record dynamics of neuronal membrane potential promises to revolutionize our understanding of neurobiology. In this study, we show that the far-red voltage sensitive fluorophore, Berkeley Red Sensor of Transmembrane potential −1, or BeRST 1, can be used to monitor neuronal membrane potential changes across dozens of neurons at a sampling rate of 500 Hz. Notably, voltage imaging with BeRST 1 can be implemented with affordable, commercially available illumination sources, optics, and detectors. BeRST 1 is well-tolerated in cultures of rat hippocampal neurons and provides exceptional optical recording fidelity, as judged by dual fluorescence imaging and patch-clamp electrophysiology. We developed a semi-automated spike-picking program to reduce user bias when calling action potentials and used this in conjunction with BeRST 1 to develop an optical spike and connectivity analysis workflow (OSCA) for high-throughput dissection of neuronal activity dynamics in development and disease. The high temporal resolution of BeRST 1 enables dissection of firing rate changes in response to acute, pharmacological interventions with commonly used inhibitors like gabazine and picrotoxin. Over longer periods of time, BeRST 1 also tracks chronic perturbations to neurons exposed to amyloid beta (Aβ^1-42^), revealing modest changes to spiking frequency but profound changes to overall network connectivity. Finally, we use OSCA to track changes in neuronal connectivity during development, providing a functional readout of network assembly. We envision that use of BeRST 1 and OSCA described here will be of use to the broad neuroscience community.

**Significance Statement:** Optical methods to visualize membrane potential dynamics provide a powerful complement to Ca^2+^ imaging, patch clamp electrophysiology, and multi-electrode array recordings. However, modern voltage imaging strategies often require complicated optics, custom-built microscopes, or genetic manipulations that are impractical outside of a subset of model organisms. Here, we describe the use of Berkeley Red Sensor of Transmembrane potential, or BeRST 1, a far-red voltage-sensitive fluorophore that can directly visualize membrane potential changes with millisecond resolution across dozens of neurons. Using only commercially available components, voltage imaging with BeRST 1 reveals profound changes in neuronal connectivity during development, exposes changes to firing rate during acute pharmacological perturbation, and illuminates substantial increases in network connectivity in response to chronic exposure to amyloid beta.

## Introduction

Rapid changes in membrane potential, or action potentials, underlie the physiology of the nervous system. Action potential firing depends on both extrinsic and intrinsic factors, many of which are neuronal cell-type specific. Brain function arises not from the behavior of isolated cells but instead from the concerted action of neuronal ensembles, making connectivity between neurons a fundamental prerequisite of neural circuits. The function of any neuronal circuit reflects the heterogeneous activity of its components. Techniques that provide high sampling can give a ‘voice’ to all neuronal types, providing a more representative overview of network function.

Studying neuronal connectivity throughout development increases our understanding of brain circuitry and function, and insights into the mechanisms of neuronal circuit construction may help shed light on how network activity breaks down in aging and disease. Indeed, network dysfunction is a hallmark of neurological diseases, including epilepsy and neurodegenerative disease. An outstanding challenge in these diseases is to define the mechanisms driving network dysfunction. Despite the importance of localizing the where and when of neuronal activity within the context of development, health, and disease, it has been difficult to capture a view of entire network behavior while maintaining cellular resolution.

Part of this challenge is due to limitations in observing and recording network activity in real time with cellular resolution. Electrophysiological techniques are often employed for direct electrical measurements. However, these methods are tedious. Simultaneous recording of multiple neurons is limited to a few neurons per experiment; therefore, information about network interplay is lost. On the other hand, macro-scale brain imaging approaches, including fMRI, positron emission tomography (PET) and EEG, resolve interactions among and between brain regions. However, these approaches cannot provide single-neuron resolution. Studies on the meso-scale, at the level of neuronal circuits but with micrometer spatial resolution, bridge the gap between micro- and macro-scale studies to investigate interactions of neurons within circuits.

Recent studies of functional interactions on the meso-scale rely on multi-electrode arrays (MEAs) which allow simultaneous measurements from tens to thousands of electrodes, or Ca^2+^ imaging, which sacrifices a direct measure of electrical activity for the ability to record from large numbers of cells.(Chang, 2015) Voltage imaging is an attractive complement to electrode-based recordings and Ca^2+^ imaging because it provides a direct readout of electrical changes while maintaining the spatial resolution of light microscopy. Voltage imaging with fluorescent indicators (Salzberg et al., 1973; Fluhler et al., 1985; Siegel and Isacoff, 1997; Sakai et al., 2001; Ataka and Pieribone, 2002; Braubach et al., 2015; Kamino, 2015) has been a long-standing goal of the scientific community; however, recent developments in both small molecule dyes, (Miller et al., 2012; Yan et al., 2012) genetically encoded voltage indicators, (Jin et al., 2012; Hochbaum et al., 2014; St-Pierre et al., 2014; Gong et al., 2015; Piatkevich et al., 2018) and combinations of the two (Abdelfattah et al., 2019; Sundukova et al., 2019; Deal et al., 2020) have prompted renewed interest in voltage imaging. Our group has been exploring voltage-sensitive fluorophores (VoltageFluors) that use photoinduced electron transfer (PeT) as a voltage sensing trigger.(Liu and Miller, 2020) VoltageFluors, possess good sensitivity, have fast response kinetics that enable single trial action potential detection, and are bright enough to be imaged with commercially available low-power LEDs and CMOS cameras. One such VoltageFluor is Berkeley Red Sensor of Transmembrane potential 1, or BeRST 1.(Huang et al., 2015)

In this manuscript, we show that voltage imaging of dissociated hippocampal neurons with BeRST 1 faithfully reports neuronal action potentials across dozens of neurons simultaneously. Along with a semi-automated action potential detection routine, BeRST 1 can rapidly interrogate neuronal excitability and connectivity changes in response to pharmacological manipulation. Finally, we characterize the effects of a pathological challenge, Aβ^1-42^, on neuronal firing rates and connectivity.

## Materials and Methods

### Cell Culture

All animal procedures were approved by the UC Berkeley Animal Care and Use Committees and conformed to the NIH Guide for the Care and Use of Laboratory Animals and the Public Health Policy.

### Rat Hippocampal Neurons

Hippocampi were dissected from embryonic day 18 Sprague Dawley rats (Charles River Laboratory) in cold sterile HBSS (zero Ca^2+^, zero Mg^2+^). All dissection products were supplied by Invitrogen, unless otherwise stated. Hippocampal tissue was treated with trypsin (2.5%) for 15 min at 37 °C. The tissue was triturated using fire polished Pasteur pipettes, in minimum essential media (MEM) supplemented with 5% fetal bovine serum (FBS; Thermo Scientific), 2% B-27, 2% 1 M D-glucose (Fisher Scientific) and 1% GlutaMax. The dissociated cells were plated onto 12 mm diameter coverslips (Fisher Scientific) pre-treated with PDL at a density of 30-40,000 cells per coverslip in MEM supplemented media (as above). Neurons were maintained at 37 °C in a humidified incubator with 5% CO_2_. At 1 day in vitro (DIV), half of the MEM supplemented media was removed and replaced with Neurobasal media containing 2% B-27 supplement and 1% GlutaMax. Functional imaging was performed on 8-15 DIV neurons to access neuronal excitability and connectivity across different stages of development.

### VoltageFluor/BeRST 1 Stocks and Cellular Loading

For all imaging experiments, BeRST 1 was diluted from a 250 μM DMSO stock solution to 0.1-1 μM in HBSS (+Ca^2+^, +Mg^2+^, -phenol red). To load cells with dye solution, the media was first removed from a coverslip and then replaced with the BeRST-HBSS solution. The dye was then allowed to load onto the cells for 20 minutes at 37 °C in a humidified incubator with 5% CO_2_. After dye loading, coverslips were removed from the incubator and placed into an Attofluor cell chamber filled with fresh HBSS for functional imaging.

### Drug Treatments

#### TTX, Gabazine, and Picrotoxin Treatments

TTX (Abcam), Gabazine (EMD Millipore), and Picrotoxin (Sigma-Aldrich) were diluted to 1 μM, 10 μM, and 50 μM respectively in HBSS (+Ca2+, +Mg2+, -phenol red). Coverslips were loaded with BeRST 1 as outlined in above. After dye loading, coverslips were placed into an Attofluor chamber filled with the Drug-HBSS solutions for functional imaging.

#### Aβ Treatments

Amyloid-beta (Aβ) peptides were purchased from AnaSpec and solubilized in a manner analogous to previous protocols (Stine et al., 2003). Briefly, lyophilized peptides were dissolved in 1,1,1,3,3,3-hexafluoro-2-propanol (HFIP) to generate a 1 mM solution. The Aβ-HFIP solution was aliquoted into PCR tubes and evaporated to dryness by placing the open PCR tubes in a chemical fume hood overnight followed by further concentration on a vacuum concentrator. The dried peptide aliquots were then stored over desiccant in glass jars at −20 °C. Before addition to the cultures, the dried peptides were re-dissolved to 1 mM in DMSO, vortexed, and sonicated for 10 minutes. Further dilutions were made with DMSO to 50-500 μM. At 8 DIV, the DMSO stocks were diluted 1:500 in fresh Neurobasal media containing 2% B-27 supplement and 1% GlutaMax (NB++). The Aβ-NB++ solutions were then added to an equal amount of conditioned media on the coverslips to give a total dilution of 1:1000 for Aβ:Media. The cultures were grown in the Aβ-NB++ solution for 7 days and loaded with dye and imaged at 15 DIV. For vehicle controls, an equal amount of DMSO containing no Aβ was added to the NB++.

### Imaging Parameters

#### Spontaneous Activity Imaging

Spontaneous activity imaging was performed on an upright AxioExaminer Z-1 (Zeiss) or an inverted Zeiss AxioObserver Z-1 (Zeiss), both equipped with a Spectra-X light engine LED light (Lumencor), and controlled with Slidebook (3i). Images were acquired using a W-Plan-Apo/1.0 NA 20x water immersion objective (Zeiss) or a Plan-Apochromat/0.8 NA 20x air objective (Zeiss). Images (2048 × 400 px^2^, pixel size: 0.325 × 0.325 μm^2^) were collected continuously on an OrcaFlash4.0 sCMOS camera (sCMOS; Hamamatsu) at a sampling rate of 0.5 kHz, with 4×4 binning, and a 633 nm LED excitation light power of 13 mW/mm^2^.

#### Image Analysis

All imaging analysis was performed using SpikeConnect, a MATLAB script developed in-house to detect action potentials from fluorescence traces associated with manually drawn regions of interest. A brief description of the SpikeConnect workflow is outlined below. This code is available from GitHub upon request. The code was added to the MATLAB path and can be run un on all versions post 2017a. The imaging data was organized in the following manner. Each coverslip was organized into individual folders which contained separate folders for each area imaged on that coverslip. Each area folder contained a brightfield image of the area and the fluorescence movies recorded from that area. All plotting and statistical analysis was performed in GraphPad.

#### Drawing Regions of Interest (ROIs) and Labeling Neurons

Individual neurons were labeled by running “selectroi_gui”, selecting a brightfield image (as a .tiff file) and importing the movies associated with the image (also as .tiff files). The frame rate parameter was set to the recording frame rate. Regions of interest (ROIs) were drawn around each neuron in the brightfield image and saved. Next, a ROI was drawn around an area of background in the first frame of the fluorescence movie associated with the brightfield image and saved.

#### Action Potential Detection

Action potentials (spikes) were detected by running “batchkmeans_gui”, selecting area folders for analysis, and using the background ROI to generate background-corrected fluorescence traces for each neuron ROI. This script then uses k-means clustering to identify possible action potentials (spikes), subthreshold events, and baseline. After the running the k-means clustering algorithm, a signal-to-noise ratio (SNR) threshold for a signal to be labeled as a spike was set by running “thresholding_gui”. This threshold was established from dual electrophysiology-optical experiments as the optical trace threshold which faithfully reproduces the spikes detected by electrophysiology. The spike data was then saved for further analysis.

#### Firing Frequency Analysis

Firing frequencies from the spike data were exported as excel files for each ROI by running “freqexport_gui”. The excel files generated contained average frequency (Hz), instantaneous frequency (Hz), and interspike interval (ms) data for each ROI along with summary columns for each of these three parameters. This data was then plotted in GraphPad for comparison.

#### Area Under the Curve (AUC) Analysis

Area under the curve (auc) data was calculated by selecting spike data and running “auc_gui”. The resulting auc data (ms) for the multi-spike averages, multi-spike sums, and whole traces was then exported as excel files and plotted in GraphPad for comparison.

#### Spike Time Tiling Coefficient (STTC) Analysis

Spike time tiling coefficients (STTC) are the correlation between a pair of spike trains. These were calculated by selecting spike data and running “sttc_gui”. The resulting STTC values for each ROI pair were then exported as an excel file and plotted in GraphPad for comparison.

#### Electrophysiology

For electrophysiological experiments, pipettes were pulled from borosilicate glass (Sutter Instruments, BF150-86-10), with a resistance of 5–8 MΩ, and were filled with an internal solution; (in mM) 115 potassium gluconate, 10 BAPTA tetrapotassium salt, 10 HEPES, 5 NaCl, 10 KCl, 2 ATP disodium salt, 0.3 GTP trisodium salt (pH 7.25, 275 mOsm). Recordings were obtained with an Axopatch 200B amplifier (Molecular Devices) at room temperature. The signals were digitized with a Digidata 1440A, sampled at 50 kHz and recorded with pCLAMP 10 software (Molecular Devices) on a PC. Fast capacitance was compensated in the on-cell configuration. For all electrophysiology experiments, recordings were only pursued if the series resistance in voltage clamp was less than 30 MΩ.

For whole-cell, current clamp recordings in hippocampal neurons, following membrane rupture, resting membrane potential was assessed and recorded at I = 0 and monitored during the data acquisition.

## Results

### Validation of optical spike detection and analysis with BeRST 1

BeRST 1 is a silicon-rhodamine-based voltage sensitive fluorescent indicator.(Huang et al., 2015) We previously reported that BeRST 1 possesses far-red to near infra-red excitation and emission profiles, excellent photostability relative to first-generation voltage sensitive indicators(Miller et al., 2012; Woodford et al., 2015)and linear fluorescence responses to changes in membrane potential. In this study, we further examined the ability of BeRST 1 to report nascent neuronal activity and connectivity in cultured rat hippocampal neurons (**Scheme 1**). In order to maximize signal-to-noise ratios (SNR) for longer recordings of spontaneous activity, we assessed combinations of light power and BeRST 1 concentration that could be tolerated before phototoxic effects were observed. We found that for the same SNR, phototoxicity was observed less when minimizing dye concentration was prioritized over minimizing light power (**Fig. 1-1**). Bath application of 500 nM BeRST 1 excited under mild illumination intensity (13 mW/mm^2^) provided stable optical recordings for >20s. Following loading, BeRST 1 localized to the plasma membranes of cell bodies and processes (**Fig. 1a,b**) with neuronal somatic regions of interest (ROIs) producing traces that show fast depolarisations that occur randomly in time.

**Figure 1.**
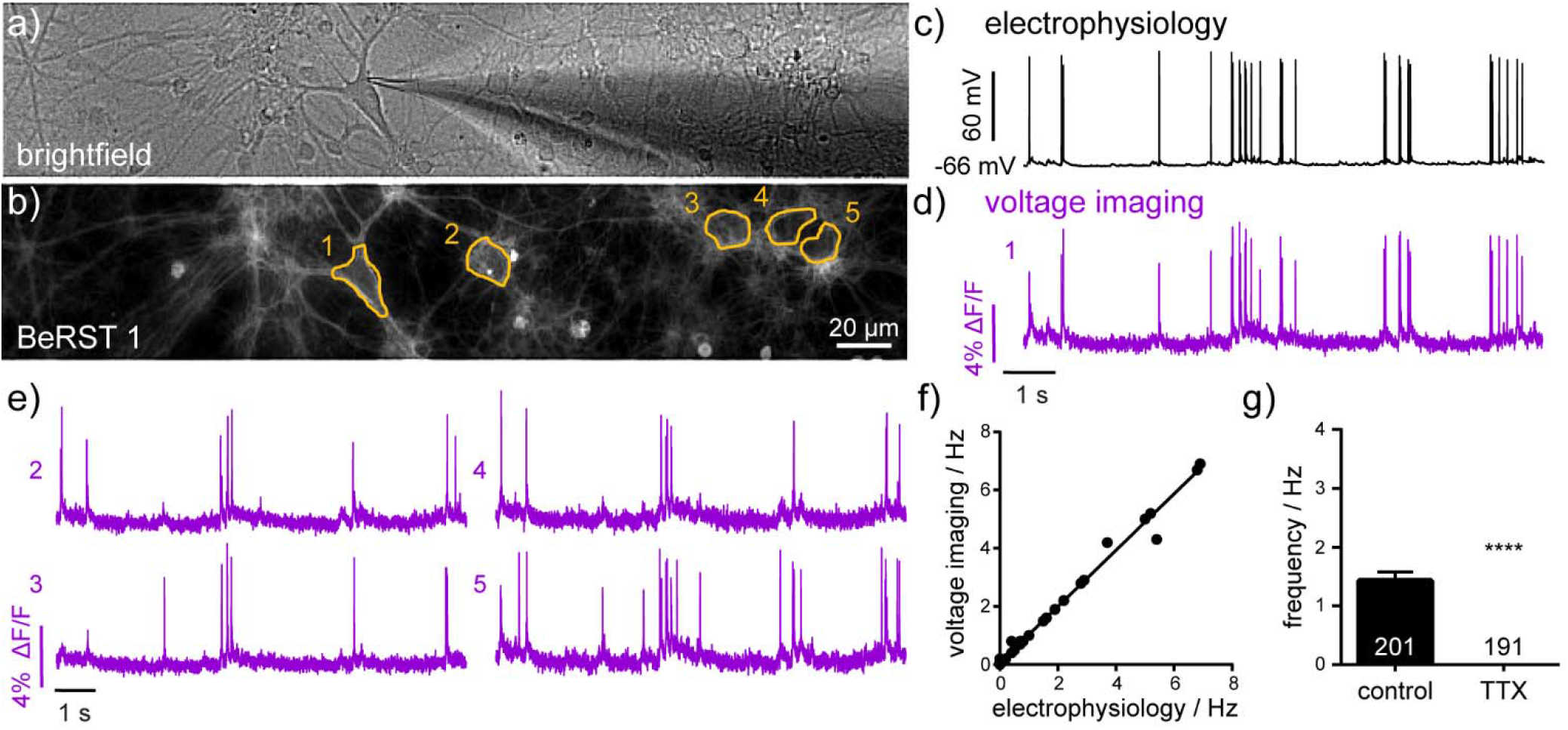
Electrophysiological characterization of voltage-sensitive fluorophore, BeRST 1. **(a)** Brightfield image of dissociated hippocampal neurons in culture showing patch-clamp targeting of a single neuron. **(b)** Fluorescence image of neurons loaded with BeRST 1 voltage-sensitive dye (500 nM). Yellow outlines indicate regions of interest (ROIs) used in **(d-e)**. Example traces from simultaneous **(c)** electrophysiological and **(d)** optical recording of spontaneous voltage fluctuations from neuron 1. **(e)** Representative BeRST 1 traces of spontaneous activity from neurons 2-5. **(f)** Action potential frequency recorded optically with BeRST 1 as a function of action potential frequency recorded electrophysiologically. Sample size, *n*, is 36 movies from 15 cells; Spearman Correlation P<0.0001, R^2^=0.99. (**g**) Firing frequency of neurons treated with 1 μM TTX compared to control sister neurons. Sample size, *n*, specified on graph are numbers of neurons, biological *n* is 3 experiments; Mann-Whitney test P<0.0001.

**Figure 1-1.**
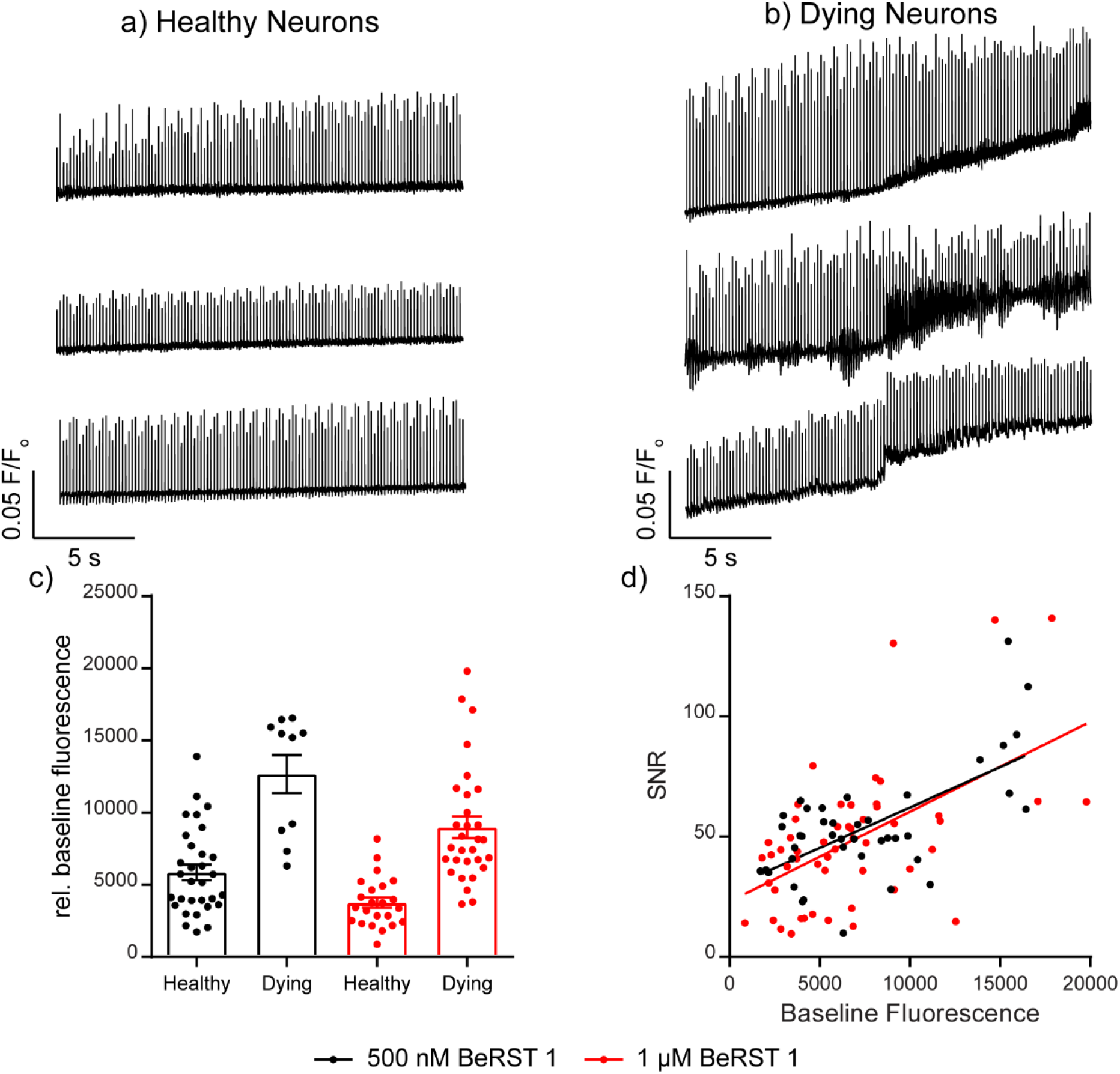
Optimization of BeRST 1 light power. Representative raw fluorescence traces normalized to baseline fluorescence values used to score neurons as **a**) healthy or **b**) dying. **c**) Plot of the baseline fluorescence values for neurons characterized as either healthy or dying at 500 nM or 1 μM BeRST 1 dye loading. **d**) Plot of signal-to-noise ratios (SNR) for action potentials from traces with varying baseline fluorescence values. The correlation between SNR and baseline fluorescence is very similar for either 500 nM or 1 μM BeRST 1 dye loading.

A challenge in recording spontaneous, as opposed to evoked activity, is spike detection. While principal/independent component analysis (PCA-ICA) analysis techniques are often applied to analyze large functional imaging datasets,(Hill et al., 2010; Frady et al., 2016) they spatially segment neurons based on activity profiles, thereby omitting silent/quiescent neurons, which are important circuit components.(Ovsepian, 2019) To address this, we developed an ROI-based semi-automated routine for action potential detection in MATLAB, which we call SpikeConnect. SpikeConnect avoids user bias by using a k-means(MacQueen, 1967; Xu and Tian, 2015) unsupervised clustering(Kiselev et al., 2019) algorithm to group intensity values into three separate bins. SpikeConnect groups putative action potentials as points within the highest intensity cluster and baseline as the lowest intensity. A signal to noise threshold is set and spike timings extracted for each action potential from every neuron recorded. We added additional modules to SpikeConnect to calculate other metrics from the fluorescence traces including the area under the curve (AUC), and the spike timing tiling co-efficient (STTC)(Cutts and Eglen, 2014) between pairs of spike trains to determine network connectivity, which will be addressed in further detail later in the manuscript.

To assess the utility of BeRST 1 voltage imaging coupled with SpikeConnect algorithms to detect somatic action potentials we compared BeRST 1 optical signals with simultaneously recorded electrophysiological traces using whole-cell patch-clamp electrophysiology (**Fig. 1c,d**). A plot of action potential frequency recorded optically and analyzed with SpikeConnect vs. frequency measured with electrophysiology and analyzed with Clampfit software reveals a near-perfect correlation between the imaging and electrode approach (**Fig. 1f**, comparisons across 36 recordings/movies from 15 different cells, R^2^= 0.99) confirming BeRST 1 imaging, in combination with SpikeConnect analysis, faithfully reports action potentials during bouts of spontaneous activity. While the signal-to-noise of the whole-cell, patch-clamp recording is superior to the optical BeRST 1 recording, BeRST 1 readily resolves action potentials and sub-threshold events (**Fig. 1c,d**) and can do so from multiple cells simultaneously (**Fig. 1e**). The small jitter in optical action potential height (**Fig. 1d**) relative to the patch-clamp recording (**Fig. 1c**) is likely a result of the lower optical sampling rate (500 Hz) relative to the electrophysiology (20 kHz). In addition, treatment of cultures with the action potential blocker, sodium channel antagonist tetrodotoxin (TTX, 1 μM, n = 191 neurons), completely abolished spontaneous action potential activity recorded optically (**Fig. 1g**).

In summary, combining BeRST 1 voltage imaging with SpikeConnect algorithms, or optical spiking and connectivity assay (OSCA), enables detection of spontaneous action potential firing with similar confidence to gold-standard electrophysiological techniques. Our voltage imaging approach dramatically increases throughput compared to patch-clamp electrophysiology. The combination of precise electrical readout of action potentials on fast temporal scales with improved throughput places OSCA in a position to evaluate changes in activity across neuronal networks following both pharmacological and pathophysiological modulations.

### Characterization of hyperactivity following pharmacological modulation

First, we wanted to test the utility of OSCA to detect pharmacological modulators which could enhance neuronal activity. Hippocampal dissociated cultures are highly connected and show spontaneous and synchronous activity in basal conditions (**Fig. 1**), and provide a powerful model system for the induction of epileptic activity (Furshpan and Potter, 1989; Vedunova et al., 2013). A simple way to achieve this is to block the activation of ionotropic GABA_A_ receptors, which generate inhibitory postsynaptic potentials (IPSCs) during synaptic transmission. This phasic inhibition is both essential in preventing overexcitation, leading to pathological states, as well as generating rhythmic activities in neuronal networks.(Farrant and Nusser, 2005) We treated 14 -17 DIV hippocampal cultures for 20 minutes with vehicle or gabazine (GBZ, 10 μM), a competitive GABA_A_R antagonist or picrotoxin (PTX, 50 μM), a non-competitive inhibitor preferentially interacting with agonist-bound GABA_A_Rs.(Newland and Cull-Candy, 1992) The high SNR and throughput of OSCA allowed us to rapidly isolate action potentials for 732 neurons. We found that treatment with either gabazine or picrotoxin results in a nearly 3-fold increase in mean firing frequency (**Fig. 2d).** Control, untreated, cultures fired at a rate of 0.9 Hz (± 0.08 Hz, S.E.M.), gabazine-treated cultures at 2.8 Hz (± 0.2, S.E.M.), and pictotoxin-treated cultures at 2.8 Hz (± 0.1, S.E.M.). Consistent with network-wide lifting of synaptic inhibition, the distribution of action potential frequencies shifts to higher frequencies (**Fig 2e**). Inhibitor-treated neurons show a decrease in the proportion of quiescent neurons, indicated by a change in Y-intercept of the cumulative frequency from approximately 47% of neurons quiescent in control cultures to 29% in gabazine- and 19% in picrotoxin-treated cultures (**Fig. 2e**).

**Figure 2.**
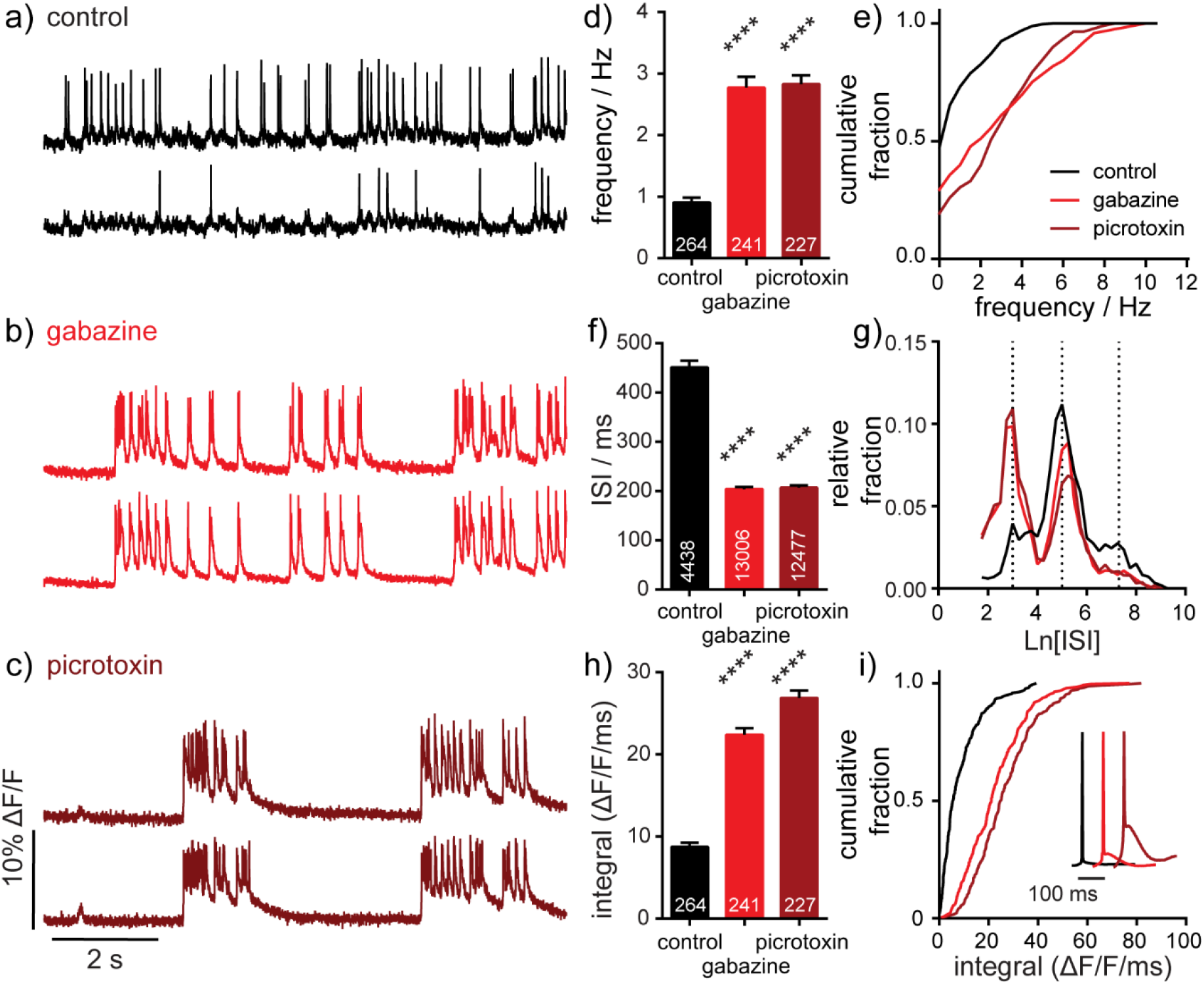
Characterization of neuronal responses to pharmacological manipulation. Representative ΔF/F voltage imaging traces of spontaneous spiking activity measured by BeRST 1 in hippocampal neurons under (**a**) control conditions or following acute administration of (**b**) 10 μM gabazine or (**c**) 50 μM picrotoxin. Traces are of 2 neurons from the same acquisition. Summarized data show plots of (**d-e**) frequency, (**f-g**) inter-spike interval (ISI), and (**h-i**) integrated area for spontaneously active neurons following acute treatment with gabazine or picrotoxin compared to sister control neurons. Data are represented as bar plots (**d,f,h**), cumulative frequency plots (**e,i**), or relative frequency distribution of natural log-transformed ISI data (**g**). Insets in panel (**i**) show mean traces scaled for amplitude. Biological *n* is 3 for gabazine and picrotoxin treatments. Values indicated on bar graphs in panels (**d**) and (**h**) indicate number of individual neurons used to determine frequency (**d**) and integrated area (**h**). Values indicated on bar graph in panel (**f**) indicate the number of pairs of consecutive action potentials used to determine ISI. Statistical tests are Kruskal-Wallis ANOVAs with multiple comparisons tests to control data. **** = p < 0.0001.

For defined imaging periods, increases in frequency correspond to decreases in inter-spike interval (ISI). We observed that treatment with GBZ and PTX significantly decreased ISI when compared to vehicle control (**Fig. 2f**). Examining the distribution of ISI values gives insight into patterns of spike timings, synchrony, and network organization. (**Fig. 2g**). Using OSCA, we find that action potentials occur within 3 discrete timing bands in control cultures: at 3 ln[ISI] (~50 Hz; 20 ms), 5 ln[ISI] (~6.6 Hz; 150 ms) and 7 ln[ISI] (~0.9 Hz; 1100 ms). In the presence of GBZ and PTX, the 3 timing bands are maintained, but inhibitor treatment evokes a redistribution in the proportion of activity within each band. High frequency (~50 Hz) firing increases 2.5 fold, balanaced by a 30-40% decrease in moderate firing (~6.6 Hz) and 50% decrease in slow firing (~0.9 Hz) frequencies (**Fig. 2g**).

GABA_A_R inhibition produces periods of sustained depolarization with APs riding on top (**Fig 2b,c**). These waveforms are highly reminiscent of paroxymal depolarizing shifts (PDS), an electrical hallmark of “ictal-like” epileptic activity (Hablitz, 1984). To quantify this activity, we calculated the response integral, or area under the curve using a custom module within SpikeConnect and found that acute gabazine and picrotoxin treatment more than doubles the ΔF/F integral (**Fig. 2h**). The cumulative frequency plot emphasizes the magnitude of the shift: 70% of control neurons show spikes that return to baseline with integrals <10 ΔF/F/ms, whereas only a small proportion do with GABA_A_R blockade (GBZ, 15%; PTX <10%, **Fig. 2i**). Interestingly, while changes in frequency and ISI are highly reproducible between GBZ and PTX (**Fig 2d-g**), we find that PTX treatment provokes larger sustained depolarizations than GBZ (**Fig. 2i**, inset, dark red vs. red). This might be due to the ability of PTX, but not GBZ, to block tonic inhibition, a mode of persistent shunting inhibition mediated by extrasynaptic GABA_A_R activity, which depresses excitatory postsynaptic potentials (EPSPs) (Farrant and Nusser, 2005). It is possible that the release of tonic inhibition by PTX leads to the generation of larger EPSPs, in turn causing greater activation of downstream effectors including voltage-dependent calcium channels thought to underlie sustained depolarization states in PDS generation.(Kubista et al., 2019)

Together, these data show that OSCA is highly amenable to the dissection of neuronal activity following global, acute network disinhibition, phenotypically capturing several hallmarks of epilepsy-like activity. In addition to robust detection of activity changes at the neuronal level, the ability to record concurrently from large numbers of neurons enabled additional insights into network-wide effects, including redistribution of spike timings. Disinhibition of a neuronal network represents a major pharmacological intervention that robustly increases neuronal activity, and while generating a robust seizurogenic model in vitro, such events are rarely observed under physiological conditions. To determine whether OSCA could offer insight into a milder network challenge, we sought to study the effects of a pathophysiological challenge by amyloid beta (Aβ) on neuronal and network activity.

### Pathophysiological modulation with Aβ^1-42^ induces neuronal hyperactivity

Network dysfunction is a common feature of neurodegenerative disease. In Alzheimer’s disease (AD), the most common neurodegenerative disease, network dysfunction, or epileptiform activity, has been observed in AD patients(Sperling et al., 2009; Huijbers et al., 2015) and mouse models(Palop et al., 2007) which simulate AD. One of the hallmarks of AD is an accumulation of amyloid beta (Aβ) plaques within the brain. When applied chronically, Aβ induces synaptic dysfunction and disrupts network connectivity.(Peña et al., 2006) To study network dysfunction in AD models a plethora of available methods for recording activity have been adopted, all with their inherent limitations, often in throughput or spike resolution (Busche et al., 2012; Verret et al., 2012; Ciccone et al., 2019a; Zott et al., 2019). As an area of vital and active research, we wanted to examine the utility of BeRST 1 voltage imaging, to dissect the impact of Aβ on neuronal network function in high detail.

Chronic exposure of developing hippocampal cultures to varying concentrations of Aβ^1-42^ (50 nM to 1 μM, **Fig. 3b,c**) for 7 days increases neuronal firing frequency in a dose-dependent manner, reaching an almost 2-fold increase following treatment with 1 μM Aβ^1-42^ (**Fig. 3d**). Interestingly, we do not observe a change in the number of quiescent neurons, which remains constant at approximately 50% across all Aβ^1-42^ treatment conditions and control (**Fig. 3e**), and suggests a mechanism where excitable cells are more active following prolonged exposure to Aβ^1-42^. This result contrasts with the network-wide increase in spiking neurons seen with acute GABA_A_R inhibitor treatment (**Fig. 2e** vs. **Fig. 3e**). The ISI for neurons treated with Aβ^1-42^ decreases relative to controls, consistent with an increase in activity upon Aβ^1-42^ treatment (**Fig. 3f**). Examination of the distribution of ln[ISI] values again reveals bands of spike timings clustered around values of 3, 5, and 7 ln[ISI] (50, 6.6, and 0.9 Hz, respectively; **Fig. 3g**). The shape of the ln[ISI] distributions in response to Aβ^1-42^ does not change significantly, indicating that the gross firing patterns in the Aβ^1-42^-treated cultures remain similar to controls. Aβ^1-42^ exposure causes a modest increase in the integrated response with no overt alterations to the shape of mean action potentials (**Fig. 3h,i**).

**Figure 3.**
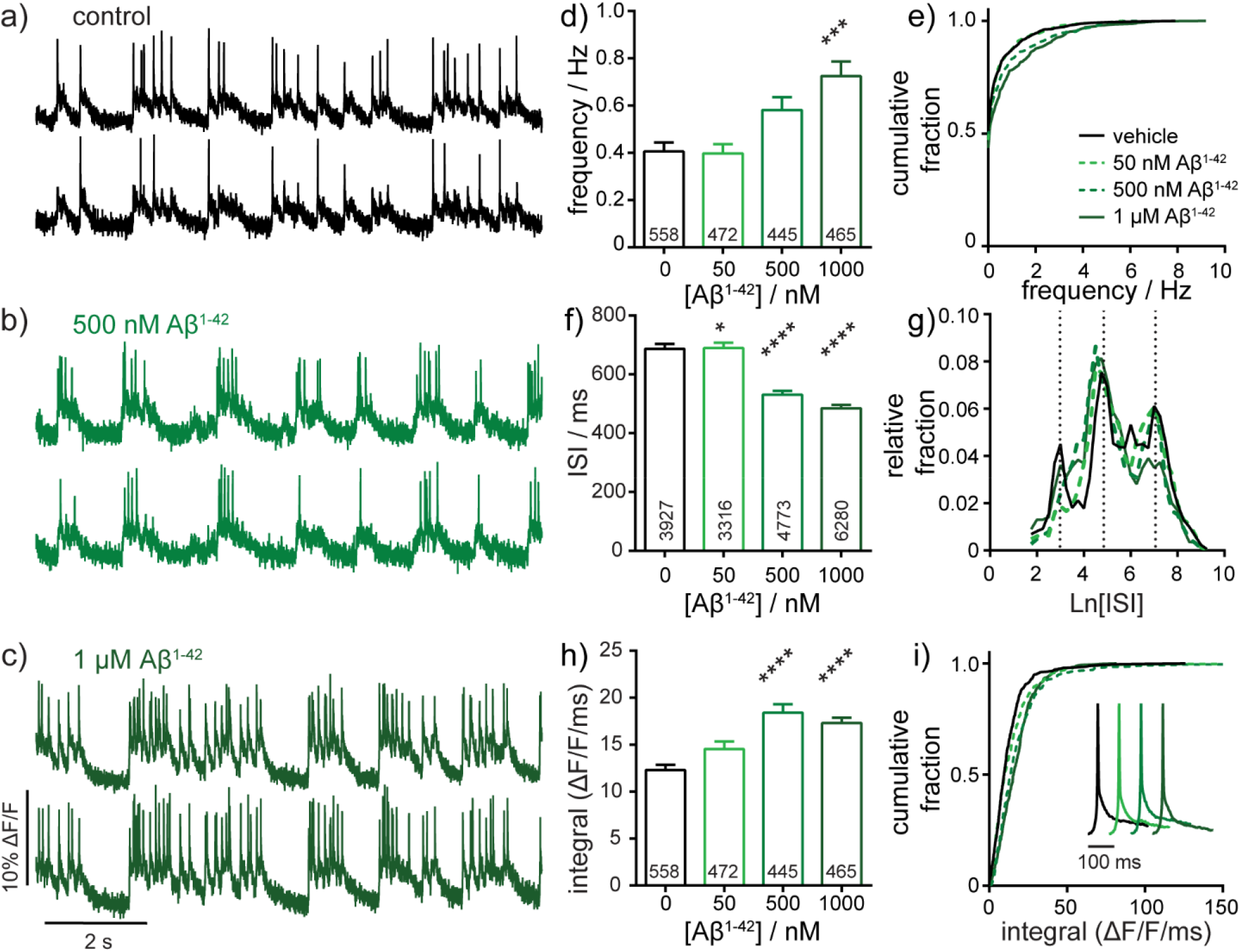
Characterization of neuronal responses upon exposure to amyloid beta 1-42 (Aβ^1-42^). Representative ΔF/F voltage imaging traces of spontaneous spiking activity in (**a**) control conditions or following chronic, 7-day administration of (**b**) 500 nM Aβ^1-42^ or (**c**) 1 μM Aβ^1-42^ peptides. Traces are of 2 neurons from the same acquisition. Summarized data show plots of (**d-e**) frequency, (**f-g**) inter-spike interval (ISI), and (**h-i**) integrated area, and for spontaneously active neurons following chronic, 7-day administration of 50 nM, 500 nM, or 1 μM Aβ^1-42^ compared to sister control neurons. Data are represented as bar plots (**d,f,h**), cumulative frequency plots (**e,i**), or relative frequency distribution of natural log transformed ISI data (**g**). Insets in panel (**i**) show mean traces scaled for amplitude. Biological *n* is 5 for Aβ^1-42^ treatments. Values indicated on bar graph in panels (**d**) and (**h**) indicate the number of neurons used to determine frequency (**d**) and integrated area (**h**). Values indicated on bar graph in panel (**f**) indicate the number of pairs of consecutive action potentials used to determine ISI. Statistical tests are Kruskal-Wallis ANOVAs with multiple comparisons tests to control data. * = p <0.05, ** = p < 0.01, *** = p < 0.001, **** = p < 0.0001.

Taken together, these data show that chronic treatment with >500 nM Aβ^1-42^ causes hyperactivity with significant increases in frequency and ΔF/F integral and decreased ISI (**Fig. 3**). The magnitude of these effects, while significant, are more subtle compared to acute GBZ and PTX (**Fig. 2**) but still readily detected by OSCA. Interestingly, hyperactivity following Aβ^1-42^ treatment manifests differently compared to GABA_A_R blockade: chronic Aβ^1-42^ treatment gives no change in fraction of quiescent neurons (**Fig. 3e**) or redistribution of ISI values (**Fig. 3g**). Chronic Aβ^1-42^ treatment, unlike acute GABA_A_R blockade, does not appear to alter the gain (or excitability) across the entire network. Rather, it appears that already-active neurons become more active, suggesting a change in underlying network connectivity. Therefore, we applied a statistical measure of functional connectivity to OSCA-detected spike trains to explore this idea further.

### Functional characterization of network formation during development

The precise timing of action potential firing between different neurons in circuits underpins the workings of the nervous systems; therefore to develop a more comprehensive picture of neural circuits, we must move away from studying single neurons in isolation, and shift towards a more holistic study of circuits.(Yuste, 2015) OSCA allows us the opportunity to infer network connectivity by evaluating temporal spiking relationships between neurons recorded simultaneously. Here, we applied a statistical method of quantifying functional connectivity, the spike-time tiling co-efficient (STTC), to the precise spike timings of clusters of neurons recorded simultaneously (10-25 neurons per field of view). Among possible descriptors of neuronal correlation,(Cutts and Eglen, 2014) STTC is attractive because it resists the influence of confounding variables, including firing rate, thus enabling a more reliable assessment of correlation for networks of heterogenous cell types. To validate the use of STTC in a voltage imaging context, we took advantage of the stereotyped structural and functional maturation of synapses in dissociated hippocampal cultures.(Dotti et al., 1988; Wagenaar et al., 2006) In this model, action potentials emerge as early as 4 days *in vitro* (DIV), but synapse formation doesn’t begin in earnest until 8-10 DIV (Basarsky et al., 1994; Grabrucker et al., 2009). Synapse formation initially leads to an excess of functional connections which over the following days are pruned in a period of synaptic remodeling to generate an efficient neuronal network with a set-point of activity (Goda and Davis, 2003; Südhof, 2018).

Using OSCA, we monitored neuronal activity in rat hippocampal neuron cultures at 8, 12, and 15 DIV, assessing the evolution of activity over the course of one week of growth and development *in vitro*. We found that overall neuronal firing frequency did not scale linearly with the developmental age of neuronal cultures (**Fig. 4a**). Rather, spiking rates peaked at 12 DIV before decreasing at 15 DIV (**Fig. 4a**). In fact, examining the firing frequency alone indicates no difference between neuronal activity at 8 and 15 DIV, with the distribution of firing frequencies perfectly overlying (**Fig 4b**, light blue vs. black). However, analysis of network STTC scores reveals robust changes (**Fig. 4c,d**). We find that STTC and network connectivity increases step-wise over the 7 day period, regardless of the underlying firing rate (**Fig. 4c,d**). Generation of synchronous activity plays central roles in fine-tuning connectivity during CNS development. We find this reflected in the ISI with spike timings occurring across a broad range of intervals at 8 DIV (**Fig. 4e,f**, light blue) before sharpening into defined bands of activity with increasing maturation at DIV 12 (**Fig. 4e,f**, blue) and DIV 15 (**Fig. 4e,f**, black). These data show empirically that during development neuronal networks modulate their excitability, neuron-neuron connectivity and network synchronicity to find a set-point of efficient activity, and further validates the use of OSCA and STTC to quantify connectivity, especially in a landscape of concurrent frequency modulations.

**Figure 4.**
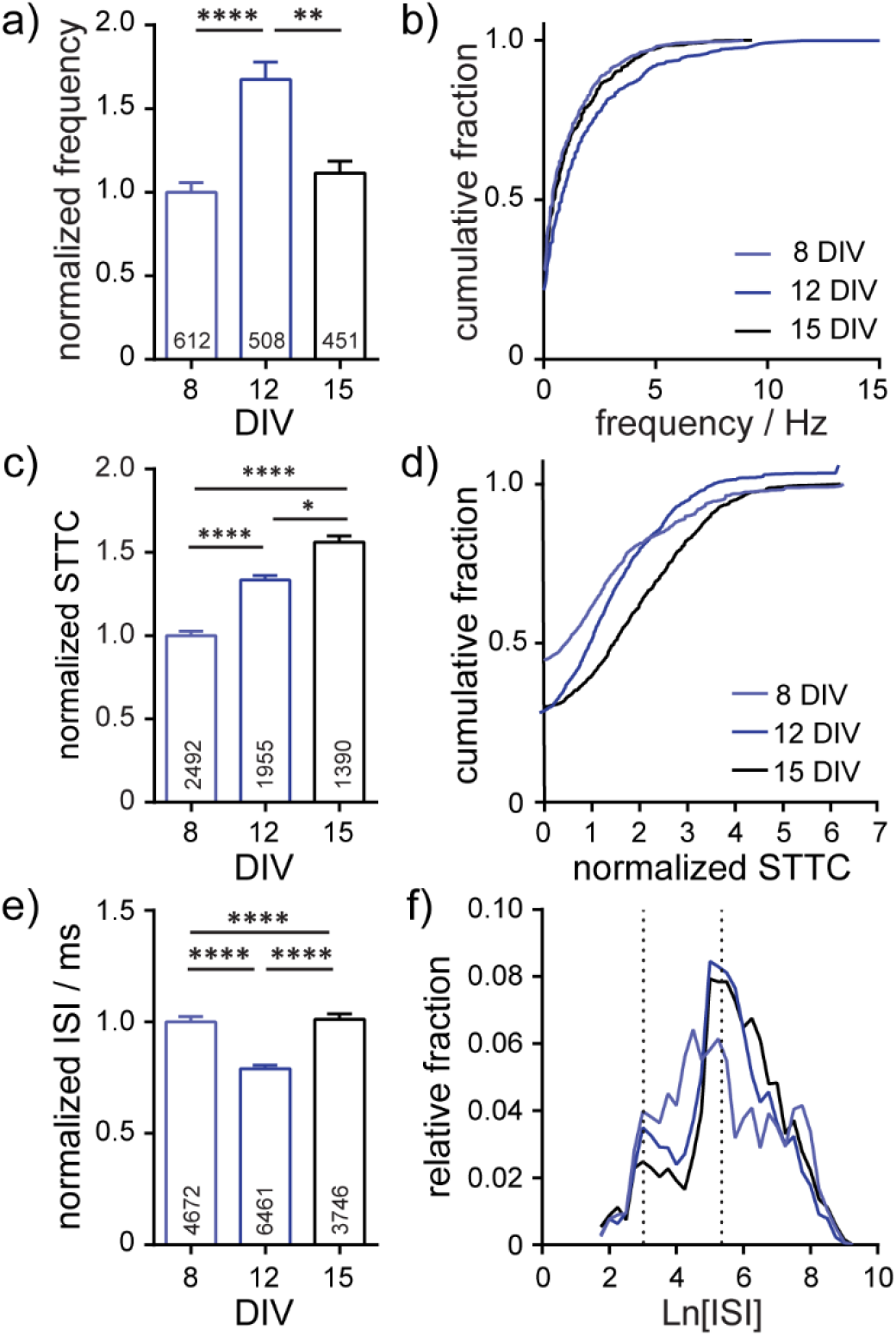
Characterization of neuronal connectivity changes over developmental stages (8, 12, and 15 *days in vitro*; DIV) and in response to pharmaceutical interventions. Summarized developmental data show plots of (**a-b**) frequency, normalized spike-time tiling co-efficient (STTC) (**c-d**), and (**e-f**) inter-spike interval (ISI) for spontaneously active neurons at the indicated stages of development. Frequency, STTC, and ISI values are normalized to the 8 DIV value per biological replicate. Frequency is summarized as a bar graph (**a**) and a cumulative frequency plot (**b**). ISI is summarized as a bar graph (**e**) and as a relative frequency distribution of the natural log transformed ISI data. Normalized STTC is summarized as bar graphs (**c**) and as cumulative frequency plots (**d**). Values on bar graphs indicate (**a**) number of neurons analyzed or (**c,e**) pairs of neurons analyzed for each condition. Data represent 3 (developmental stages) biological replicates. Statistical tests are Kruskal-Wallis ANOVAs with multiple comparisons tests to all groups (**a,c,e**). * = p <0.05, ** = p < 0.01, *** = p < 0.001, **** = p < 0.0001.

### Chronic exposure to Aβ^1-42^ induces changes to neuronal connectivity

To determine whether perturbations to neuronal networks, GABA_A_R blockade and Aβ^1-42^ treatment, would manifest changes in neuronal correlation, we analyzed network STTC. As expected, acute network-wide disinhibition with GBZ and PTX (as in **Fig. 2**) increased network synchrony: STTC values increase compared to control cultures (**Fig. 5a,b**). This is consistent with releasing the inhibitory brake on the system so that multiple neurons downstream of excitatory synaptic drivers are more likely to fire and are therefore more co-ordinated. However, the observed increase in STTC for GABA_A_R blockade is relatively small (20-30% increase) compared to the nearly 300% increase in firing frequency observed for the same treatment (**Fig. 2**). This indicates that while correlation between spike trains does increase, changes in network inter-connectivity are not a key contributor to the frequency changes under acute, global disinhibition.

**Figure 5.**
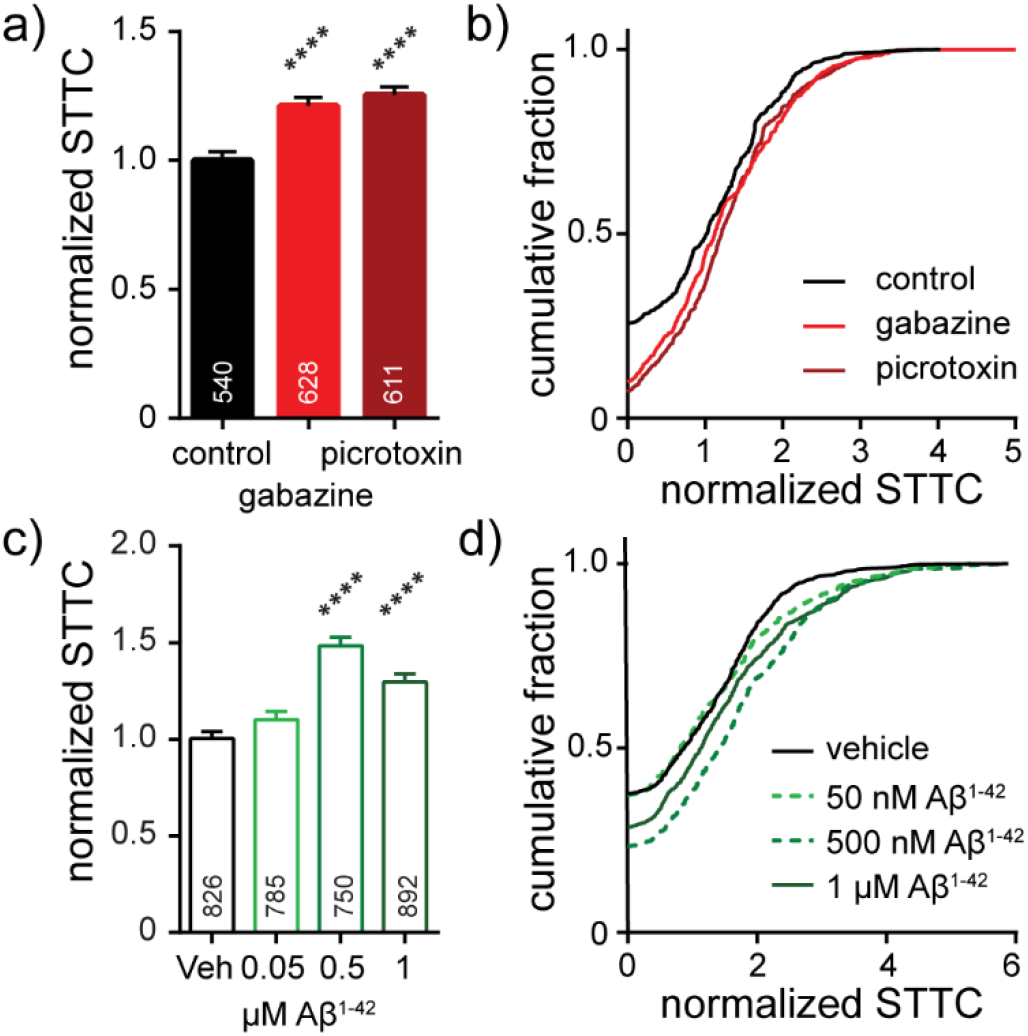
Characterization of neuronal connectivity as measured by normalized STTC for cultures (**a-b**) acutely treated with gabazine (10 μM) or picrotoxin (50 μM) or (**c-d**) following chronic, 7-day exposure to 50 nM Aβ^1-42^, 500 nM Aβ^1-42^ or 1 μM Aβ^1-42^. Normalized STTC is summarized as bar graphs (**a,c**) and as cumulative frequency plots (**b,d**). Values on bar graphs indicate pairs of neurons analyzed for each condition. Data represent 3 (gabazine/picrotoxin) or 5 (Aβ^1-42^) biological replicates. Action potential frequency and ISI for these pharmaceutical interventions are compared in **Figure 2, 3**. Statistical tests are Kruskal-Wallis ANOVAs with multiple comparisons to control (**a,c**). * = p <0.05, ** = p < 0.01, *** = p < 0.001, **** = p < 0.0001.

For chronic treatments with Aβ^1-42^ (**Fig. 3**), we also observed increases in network synchrony. Low concentrations of Aβ^1-42^ (50 nM) have little effect on either firing rate (**Fig. 3d**) or relative STTC (**Fig. 5c,d**). However, intermediate concentrations of Aβ^1-42^ (500 nM) produce a nearly 50% increase in STTC and a 25% increase at 1 μM (**Fig. 5c,d**). Interestingly, compared to treatment with either GBZ or PTZ (**Fig. 2**), Aβ^1-42^ exhibits milder effects on firing frequency, integral, and ISI (**Fig. 3**) but larger changes to the STTC value relative to controls (25 to 50% increase in STTC, **Fig. 5c,d**). Together with frequency data showing no change to the number of unresponsive neurons (**Fig. 3e**), the increase observed in STTC points to a mechanism where active neurons, under chronic exposure to Aβ^1-42^, increase the number or strength of existing connections in downstream neurons.

## Discussion

In this manuscript, we demonstrate the utility of OSCA, a voltage-sensitive fluorophore-based imaging and analysis approach in dissociated hippocampal neurons, to 1) provide high fidelity tracking of action potentials across large numbers of neurons, 2) quantify hyperactivity following pharmacological and pathophysiological network modifications, 3) validate a statistical measure of functional connectivity (Spike-Time Tiling Co-efficient; STTC) and 4) apply STTC to understand network organization in pharmacologically and pathophysiologically modified circuits.

### OSCA reliably reports activity within neuronal networks

In this study, we show that OSCA delivers detailed information on action potential firing frequency with the same resolution as the gold-standard technique for this metric, patch clamp electrophysiology (**Fig. 1**). Moreover, we demonstrate that OSCA can robustly detect network silencing (**Fig. 1**) and neuronal hyperactivity driven by either pharmacological (**Fig. 2**) or pathological agents (**Fig. 3**). However, where patch clamp is low-throughput and limited in the number of neurons recorded, OSCA is advantageous as it enables the recording of firing frequency and ISI in larger numbers of neurons simultaneously. The ability to simultaneously record from multiple cells enables OSCA to minimize cell-type bias and to gain holistic overviews of heterogenous hippocampal networks *in vitro* by examining activity distributions. For example, in developing cultures (8 DIV), we found that spike intervals were distributed across the whole range of resolvable frequencies (0-150+ Hz), but when mature, at >12 DIV, active neurons settled into 3 main bands of firing frequency centered around: 3 ln[ISI] (~ 50 Hz), 5 ln[ISI] (~6.6 Hz) and 7 ln[ISI] (~0.9 Hz) (**Fig. 4f**). Although the number of active neurons within each frequency band was redistributed after pharmacological disinhibition of the network (**Fig. 2**), the maintenance of the three activity bands suggests that these patterns of neuronal activity are not acutely controlled by inhibition.

### OSCA enables assessment of functional connectivity within networks

Neuronal connectivity in dissociated cultures cannot easily be assessed by patch clamp electrophysiology. Instead, neuronal connectivity is often assessed using MEA which, while powerful at describing dynamic states, often use global activity such as bursts or activity complexity occurring in a proportion of electrodes simultaneously(Hyvärinen et al., 2019) because ascribing single spikes to single neurons is challenging. Using an optical approach, spatial resolution is vastly improved, enabling us to confidently attribute activity signatures to individual neurons. After recording from multiple cells simultaneously, we quantified the connectedness of the underlying network using STTC, a statistical measure (STTC) of functional connectivity. As validation of this metric, we observed clear stepwise increases in functional connectivity as networks matured *in vitro*, undergoing organization and synchronization coincident with synapse formation and refinement (**Fig. 4**). These data are also consistent with MEA studies of development in cultured neurons (Wagenaar et al., 2006). After pharmacological disinhibition of the network with GABA_A_R blockers, we observed a further increase in STTC over control conditions representing an increase in network synchrony (**Fig. 4**). Together these data show that STTC is a useful readout for defining network connectivity in voltage imaging.

### Aβ^1-42^ challenge enhances neuronal firing frequency and connectivity

Understanding of the role of Aβ in modulating neuronal function is constantly evolving. Models with elevated levels of Aβ have described synapse loss, hypoactivity, and neuronal death coinciding with cognitive decline.(Shankar and Walsh, 2009) However, it is becoming increasingly evident that hyperactivity characterized by epileptiform activity is a hallmark of pre-clinical and early stage AD (Sperling et al., 2009; Huijbers et al., 2015) and is also observed in mouse models simulating AD (Palop et al., 2007). Using OSCA, we show that chronic (7 day) treatment of hippocampal cultures with > 500 nM Aβ^1-42^ peptide resulted in increased neuronal firing frequency (**Fig. 3**) and an increase in network correlation as measured by STTC (**Fig. 4**). These data link exposure to Aβ^1-42^ with neuronal hyperactivity, consistent with Ca^2+^ imaging data (Busche et al., 2012). Several mechanisms have been proposed to underlie Aβ-mediated neuronal hyperactivity, including impaired glutamate reuptake (Zott et al., 2019), reduced function of interneurons (Palop et al., 2007; Palop and Mucke, 2016), changes in neurotransmitter release probability(Abramov et al., 2009; Wang et al., 2017) and intrinsic excitability (Ciccone et al., 2019b). However, without a detailed mechanistic study, it is difficult to extrapolate effects on specific intrinsic or synaptic properties to the working of microcircuits. Here, we uncovered that relatively modest increases in firing frequency following Aβ^1-42^ treatment were accompanied by robust increases in STTC, suggesting a network that is more organized with higher levels of synchrony (**Fig. 3–4**). While the role of hyperactivity in AD is still to be fully elucidated, it may play compensatory and/or neuroprotective roles within the network (Elman et al., 2014).

### Looking to the future: a role for OSCA as a complementary screening technology

Voltage imaging methods have improved over the last decade, beginning to address problems of with poor signals, slow temporal kinetics, and cytotoxicity. BeRST 1 combines all these improvements with ease of use and widespread applicability: optical voltage recordings can be made using commercially-available, off-the-shelf imaging equipment and without the need for genetic modifications to the sample. In particular, voltage imaging with BeRST 1 is especially amenable to screening large numbers of neurons as staining intensities and therefore signal to noise ratios are relatively uniform across many cells, simplifying action potential detection. This is in contrast to genetic methods where expression levels can vary widely between cells leading to differences in the quality of recordings between cells.

The ability of OSCA to robustly detect changes in network activity profiles, coupled with the flexibility of utility in neuronal culture preparations from any host species or preparation, especially those where genetic manipulation is impossible, cumbersome, or expensive, such as patient-derived human induced pluripotent stem cell-derived neurons (Williams et al., 2019), make this technique directly amenable for high-throughput screening. In recent years there has been a resurgence in phenotypic screening for discovering new drug candidates, drug targets and for neurotoxicological testing, with a focus on *in vitro* models that translate to *in vivo* and ultimately the clinic (Moffat et al., 2017). The VoltageFluor family of dyes have shown utility in human iPSC derived neurons, a model with great disease modelling potential. Pairing OSCA with human iPSC-derived disease models will be a powerful avenue for future research, generating large functional datasets with greater translatability to humans. Similarly, our understanding of neuroscience has been limited by the lack of available tools to dissect neuronal function on the microcircuits level. Due to the complexity of even in vitro circuits it is difficult to extrapolate findings in functional changes to synaptic transmission or action potential firing to the network level. We envision that OSCA will enable a variety of novel functional studies of neuronal circuits on the meso-scale.

## Acknowledgements

We acknowledge support from the NIH (NS098088). BKR was supported in part by an NIH Training Grant (T32GM066698).

---

**Scheme 1.**
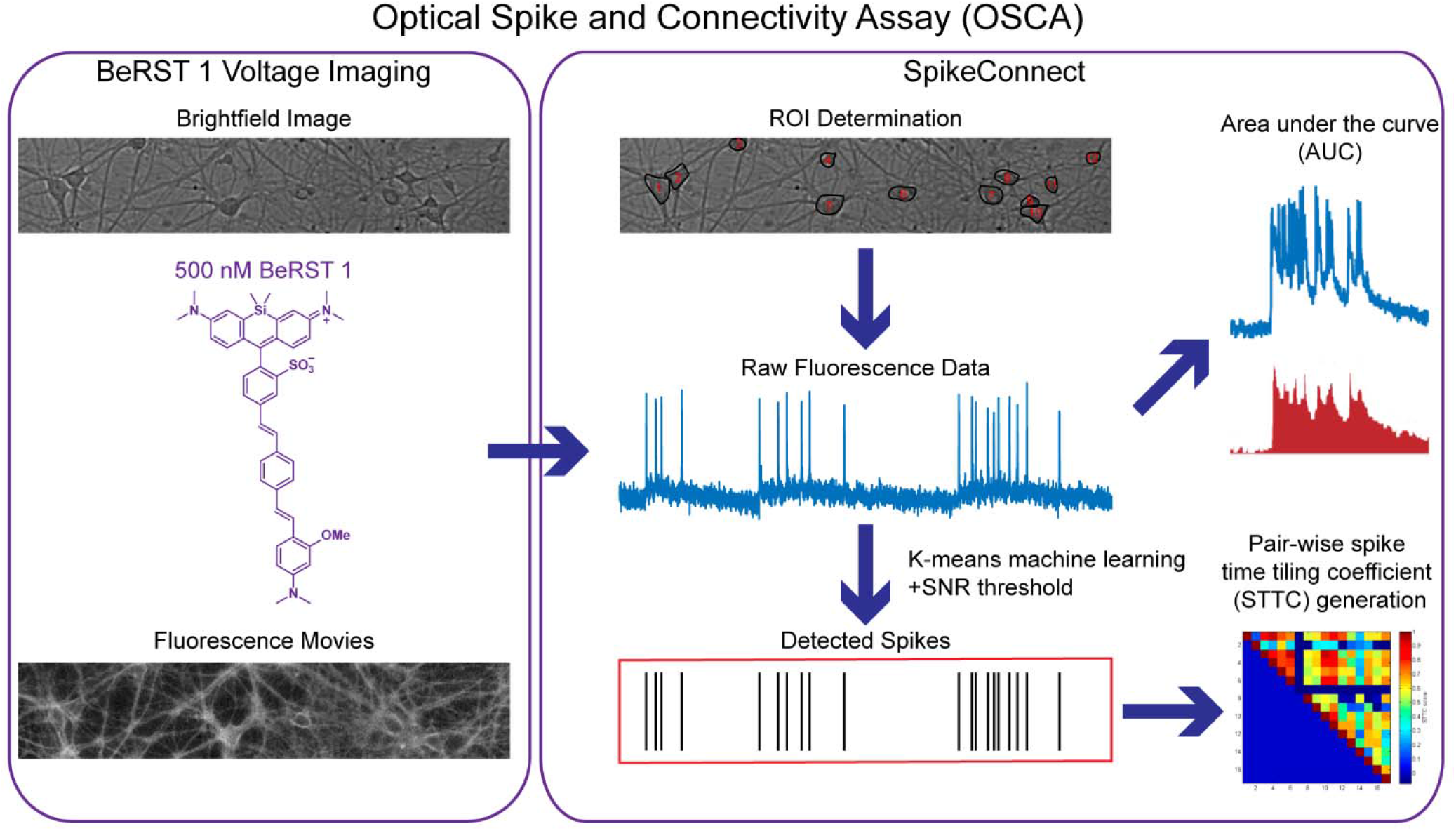
Outline of the Optical Spike and Connectivity Assay (OSCA). Using off the shelf microscope components, brightfield images and fluorescence movies are obtained from a 650 μm × 120 μm field of neurons stained with 500 nM BeRST 1. This voltage imaging data is then analyzed by SpikeConnect, a MATLAB script developed specifically for OSCA. Using SpikeConnect, regions of interest (ROIs) containing the soma of neurons are selected and the corresponding fluorescence traces for these ROIs are extracted. A k-means clustering algorithm and signal-to-noise threshold are then used to identify action potentials (spikes) from the fluorescence traces. Area under the curve can also be determined for the fluorescence traces. The spiking data for pairs of neurons can be further analyzed to determine spike time tiling coefficients (STTCs) which are statistical measures of the functional connectivity of neurons.

